# Additive genes contribute to yield heterosis in rice

**DOI:** 10.64898/2026.09.14.750853

**Authors:** Zhiwu Dan, Yunping Chen, Tianshun Zhou, Dingyang Yuan, Wenchao Huang

## Abstract

The molecular mechanisms underlying heterosis remain unresolved, largely owing to the scarcity of accurately identified heterosis-associated genes. Here, we systematically investigate the inheritance patterns of three yield heterosis-related genes in rice, analyzing their effects in both homozygous and heterozygous genetic backgrounds. We find that heterozygosity at individual loci predominantly produces additive, rather than overdominant, effects in homozygous genetic backgrounds. Nevertheless, heterozygous states of these genes generate yield heterosis in heterozygous genetic backgrounds, supporting the overdominance model. Our findings confirm that heterozygous additive genes are key contributors to yield heterosis in rice.

---

Dear Editor,

The genetic bases of heterosis, a phenomenon crucial for enhancing crop yield, remain incompletely understood. Shull’s early overdominance model proposed that heterozygosity of “genetic elements”—currently interpreted as genes—drives heterosis (Shull, 1908). East later supported the model and further suggested that heterosis arises from the cumulative action of nondefective alleles (East, 1936). Subsequently, Williams theorized that heterosis of compound traits like yield may result from interactions among additive genes, each contributing mid-parent values in F_1_ hybrids (Williams, 1959). However, because of the scarcity of functionally validated heterosis-associated genes, these models or theories have seldom been rigorously tested, and the molecular underpinnings of heterosis remain without consensus.

Crop yield is a compound trait integrating multiple components like grain number, grain weight, panicle/ear number, and seed setting rate, implying a polygenic nature of yield heterosis. A few recently identified genes for heterosis of key agronomic traits over the past three years has provided an unprecedented opportunity for in depth understanding of the genetic architecture of crop heterosis at the single-gene resolution (Sun et al., 2023; Wang et al., 2024; Dan et al., 2025; Zhang et al., 2025). Here, we analyzed the contributions of three heterosis-associated genes—*OsMADS1* (Wang et al., 2024), *OsBZR1* (Dan et al., 2025), and *Ghd8* (Sun et al., 2023)—to yield heterosis through integrating classical genetic models and inheritance patterns in rice.

Splice-site deletions in the seventh intron and eighth exon of *OsMADS1* (e.g., *OsMADS1*^*GW3p6*^, *OsMADS1*^*lgy3*^, *OsMADS1*^*-4bp*^, or *OsMADS1*^*-27bp*^), produce functional alleles that enhance yield, thousand-grain weight, grain length, and grain quality compared to the wild-types in both *indica* and *japonica* varieties (Liu et al., 2018; Wang et al., 2024). To determine the inheritance patterns of heterozygous *OsMADS1* in homozygous genetic backgrounds, phenotypes were first compared between mutated F_1_ hybrids and their parents (*indica* variety Fuhui676 and its mutants). Results indicated that heterozygosity for a single gene, *OsMADS1* (*OsMADS1_osmads1*: *OsMADS1_OsMADS1*^*-4bp*^ or *OsMADS1_OsMADS1*^*-27bp*^), conferred additive effects for thousand-grain weight and grain quality, partially dominant effect for grain length, and additive/dominant effects for grain yield (Figure 1A). Then, phenotypic values of F_6_ individuals derived from an F_5_ recombinant inbred line (RIL) of Guangzhan63-4S (*indica*) and Fuhui676—specifically, RIL79, which is heterozygous at *OsMADS1* but homozygous elsewhere—were used to compare homozygous and heterozygous genotypes of *OsMADS1* (Wang et al., 2019). Consistent with the above results of F_1_ hybrids, heterozygous *OsMADS1* (*OsMADS1_OsMADS1*^*OsGW3p6*^) exhibited near-additive effects for both grain length and thousand-grain weight (Figure 1B), highlighting the important roles of additive effects on heterosis for yield and its component traits. Notably, using another *indica* variety HJX74 as the recurrent parent, BC_4_F_2_ (backcrossing four times, BC_4_) individuals carrying heterozygous *OsMADS1* (*OsMADS1_OsMADS1*^*lgy3*^) from a *japonica*-type RIL (RIL186 derived from *japonica* variety Wuyunjing7 and American *japonica* variety L-204) also displayed additive effects for grain length, thousand-grain weight, and yield per plant (Liu et al., 2018). Furthermore, heterozygosity for single *bZIP29* produced additive effects on both its expression levels and plant/ear height in hybrid maize (Zhang et al., 2025). Collectively, these convergent findings underscore the fundamental roles of additive effects conferred by individual genes on yield heterosis in homozygous genetic backgrounds.

**Figure 1.**
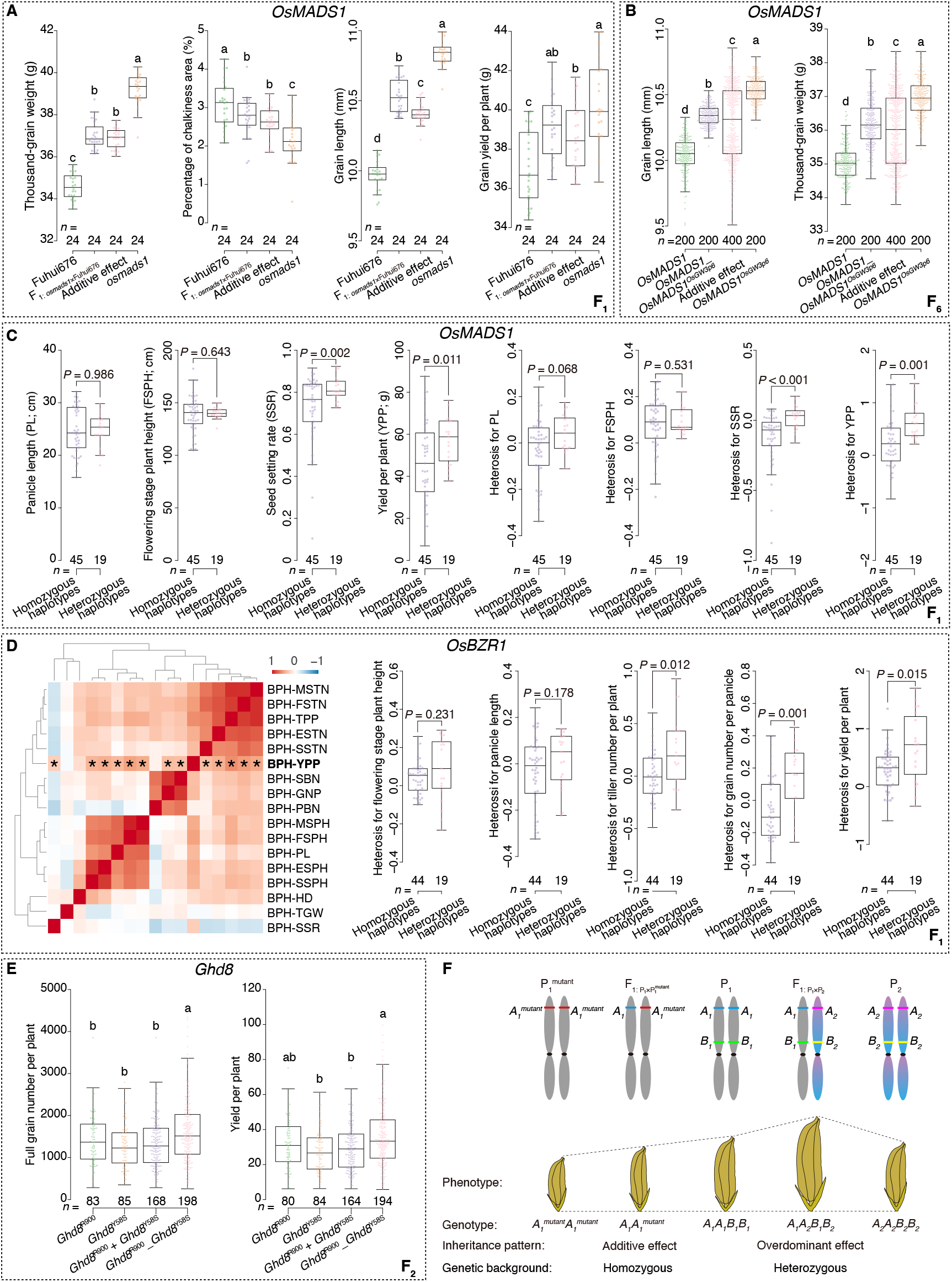
Contributions of heterosis-associated genes to phenotypes and heterosis across homozygous versus heterozygous genetic backgrounds in rice. (**A**) Phenotypic values of Fuhui676, *OsMADS1* splice-site mutants (carrying −4 bp or −27 bp deletions), their F_1_ hybrids, and the calculated additive effect (average values of Fuhui676 and its mutants). (**B**) Grain length and grain weight of F_6_ individuals derived from an F_5_ recombinant inbred line heterozygous for *OsMADS1. n* = number of plants in **A-B.(C)** Phenotypic values and better-parent heterosis of F_1_ hybrids with homozygous (haplotype-a, haplotype-b) or heterozygous (haplotype-a_haplotype-c, haplotype-b_haplotype-c) haplotypes of *OsMADS1*. Based on previous results (Dan et al. 2025), haplotype-c is the advantageous haplotype for yield heterosis, and its combinations with haplotype-a or haplotype-b are treated as heterozygous haplotypes. Haplotype-c includes inbred line Qianlijing. (**D**) Correlations of heterosis and heterosis of F_1_ hybrids with homozygous (haplotype-*indica*, haplotype-*japonica*) or heterozygous (haplotype-*indica*_haplotype-JR2, haplotype-*japonica*_haplotype-JR2) haplotypes of *OsBZR1*. Haplotype-JR2 is the advantageous haplotype for yield heterosis and includes inbred line JR2. Asterisks represent significant Pearson correlations (*P* < 0.05). *n* = number of F_1_ hybrids in **C**-**D**. (**E**) Phenotypic values of Y900 F_2_ individuals with homozygous *Ghd8*, their averages, and the heterozygous genotypes. *n* = number of F_2_ individuals. (**F**) A model for the contributions of heterosis-associated genes to yield across homozygous and heterozygous genetic backgrounds. P_1_^mutant^ represents the mutated line of Parent_1 (P1), while P2 is a genetically distinct line. *A*_*1*_ and *A*_*1*_^mutant^ represent the wild-type and mutant alleles of gene *A*, respectively. *A*_*2*_ and *B*_*2*_ are alleles of *A*_*1*_ and *B*_*1*_, respectively. Analysis of variance with fixed factors in simple linear models was performed to compare the differences in **A, B, E**, with pairwise comparisons in post hoc test. Independent samples *t-*test or Welch’s *t*-test were performed depending on equality of variances in **C**-**D**, with *P* values indicated.

Leveraging the three previously established haplotypes of *OsMADS1* (Dan et al., 2025), of which haplotype-c is characterized as favorable for yield heterosis, its effects on heterosis of 17 agronomic traits was investigated in heterozygous genetic backgrounds. This was achieved through crosses involving 12 representative inbred lines, except for *indica* varieties Mianhui725 and 610234, most of which have no overlapping parentage with Fuhui676 or Guangzhan63-4S (https://www.ricedata.cn). The resulting F_1_ hybrids, harboring either homozygous (haplotype-a or haplotype-b) or heterozygous (haplotype-a_haplotype-c or haplotype-b_haplotype-c) haplotypes, showed varied heterosis (Figure S1). Yield heterosis was significantly correlated with heterosis of traits like seed setting rate, panicle length, and plant height. Notably, the hybrids with heterozygous haplotypes had significantly higher phenotypic values and stronger heterosis than the pooled homozygous ones for both seed setting rate and yield per plant, while were without significant difference for panicle length and plant height (Figure 1C and Figure S2). In contrast, correlation analysis of *OsBZR1* revealed that yield heterosis was significantly correlated with traits including tiller number, grain number, and panicle length (Figure 1D and Figure S3). Despite these differences in correlated traits, hybrids with heterozygous haplotypes of *OsBZR1* also exhibited superior phenotypic values and enhanced heterosis for yield and yield-components (Figure 1D and Figure S4), confirming yield advantages conferred by heterozygous genes in heterozygous genetic backgrounds.

*Ghd8* contributes to yield heterosis in the super-hybrid rice Y Liangyou 900 (Y900), derived from R900 and Y58S (Sun et al., 2023), with Y58S sharing parentage with Mianhui725. *Ghd8* knockout lines of R900 showed reduced yield, plant height, and accelerated heading, and crossing these lines to Y58S produced F_1_ hybrids with lower yield compared to Y900. To study the contribution of heterozygous *Ghd8* to hybrid phenotypes in heterozygous genetic backgrounds, 12 traits were analyzed in F_2_ individuals of Y900. Grain yield was significantly correlated with traits such as grain number and plant height (Figure S5). Notably, F_2_ individuals heterozygous at *Ghd8* (*Ghd8*^R900^_*Ghd8*^Y58S^) outperformed homozygous types (e.g., overdominant effects) or their pooled averages for yield, grain number, and plant height (Figure 1D). Thus, in contrast to the additive effects observed in homozygous genetic backgrounds, heterozygous *OsMADS1, OsBZR1*, and *Ghd8* each positively contributed to yield heterosis in heterozygous genetic backgrounds.

At a systems level, we have demonstrated that additive effects of gene expression and metabolite levels are among the predominant inheritance patterns for yield heterosis in rice (Dan et al., 2025; Dan et al., 2026), corroborating the contributions of additive effects from heterosis-associated genes at the genomic level. We here proposed a simplified model to display the contributions of heterosis-associated genes to yield across different genetic backgrounds. In a homozygous background, a mutation in gene *A* (e.g., *A*_*1*_^mutant^) that reduces yield in Parent_1 (P_1_) produces an additive effect in the F_1_ hybrid, which carries the heterozygous gene *A* (e.g., *A*_*1*_*A*_*1*_^mutant^) (Figure 1F). However, when *A*_*1*_ and *A*_*2*_ are derived from P_1_ and P_2_, respectively, heterozygosity of gene *A* (e.g., *A*_*1*_*A*_*2*_) confers an overdominant effect on yield in a heterozygous background (e.g., *A*_*1*_*A*_*2*_*B*_*1*_*B*_*2*_), resulting in positive yield heterosis (Figure 1F).

Contrasting with our findings, yield heterosis in tomato has been attributed to a single heterosis-associated gene, *SINGLE FLOWER TRUSS* (*SFT*) (Krieger et al., 2010). While this example fueled interest in utilizing single-gene yield heterosis, we concur with a subsequent study suggesting that the observed yield advantage may reflect outcomes of specific genotype-by-environment interactions—for instance, enough flowering timing in F_1_ hybrids relative to *sft* mutants (Shen et al., 2022). Under favorable growth conditions such as adequate temperature and sunlight, the *sft* mutants flower later and have a prolonged growth period, potentially yielding the highest fruit weight when fruits reach full maturity. Finally, in a homozygous genetic background, the heterozygous *SFT*_*sft* may exhibit additive rather than overdominant effect on fruit yield.

In summary, our analyses reveal that heterozygous heterosis-associated genes contribute to yield heterosis in rice, with individual genes predominantly acting in an additive manner in homozygous genetic backgrounds. Functional investigation of multiple heterosis-associated genes in the same homozygous genetic backgrounds (e.g., double or triple mutations or overexpression of *OsMADS1, OsBZR1*, and *Ghd8*) should deepen our understanding of the genetic mechanisms of yield heterosis. And systematic identification of heterosis-associated genes for all yield components and elucidating their interactions will further provide a comprehensive genetic landscape. Finally, as Williams posited more than 60 years ago (Williams, 1959), reciprocal inequality in parental contributions—either in gene expression or component traits—can generate yield heterosis in F_1_ hybrids, a principle that may inform parental combinations in hybrid breeding programs.

## Methods

Phenotypic and genotypic data of parental lines, mutants, F_1_ hybrids, F_2_ individuals, F_6_ recombinant inbred lines are obtained from previously published studies (Liu et al., 2018; Sun et al., 2023; Wang et al., 2024; Dan et al., 2025). The statistical analyses were implemented using these data also provided in Supplementary Table 1. To compare the differences among groups, analysis of variance (ANOVA) with fixed factors in simple linear models was performed, adopting pairwise comparisons in post hoc test. For two-group comparison, independent samples *t*-test or Welch’s *t*-test were conducted based on equality of variances, with *P* values showing to indicate the differences.

## Acknowledgements

We thank Mr. Caibin Zhu for the rice research fund assistance.

## Author contributions

Z.W.D. conceived and designed the study. Z.W.D., Y.P.C. and T.S.Z. performed the analyses. Z.W.D., Y.P.C., D.Y.Y. and W.C.H wrote the manuscript.

## Data availability

Phenotypic, genotypic, and haplotype data were obtained from four previously published studies (Liu et al., 2018; Sun et al., 2023; Wang et al., 2024; Dan et al., 2025) and provided in Table S1.

