## Supplementary figures 1-5 for "Additive genes contribute to yield heterosis in rice"

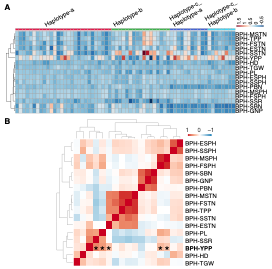

**Figure S1** Better-parent heterosis of 17 agronomic traits in 64 F_1_ hybrids with homozygous (haplotype-a, haplotype-b) or heterozygous (haplotype-a_haplotype-c, haplotype-b_haplotype-c) haplotypes of *OsMADS1*. (**A**) Heatmap for better-parent heterosis of 17 traits. (**B**) Heatmap for Pearson correlations among heterosis of 17 traits. Asterisks represent significant correlations between yield per plant and the other traits (*P* < 0.05). Abbreviations: better-parent heterosis = BPH, seedling stage plant height = SSPH, elongation stage plant height = ESPH, flowering stage plant height = FSPH, maturation stage plant height = MSPH, primary branch number = PBN, secondary branch number = SBN, grain number per panicle = GNP, seedling stage tiller number = SSTN, elongation stage tiller number = ESTN, flowering stage tiller number = FSTN, maturation stage tiller number = MSTN, tiller number per plant = TPP, panicle length = PL, seed setting rate = SSR, yield per plant = YPP, heading date = HD, and thousand grain weight = TGW.

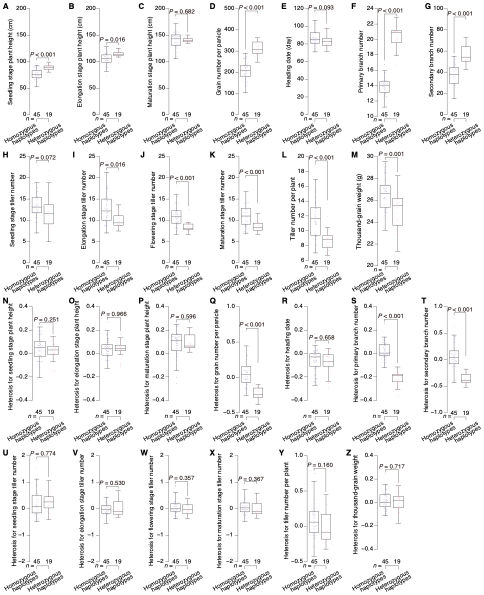

**Figure S2** Phenotypic values and better-parent heterosis in 64 F_1_ hybrids with homozygous (haplotype-a, haplotype-b) or heterozygous (haplotype-a_haplotype-c, haplotype-b_haplotype-c) haplotypes of *OsMADS1*. Panels (**A-M**) show phenotypic values for seedling stage plant height (**A**), elongation stage plant height (**B**), maturation stage plant height (**C**), grain number per panicle (**D**), heading date (**E**), primary branch number (**F**), secondary branch number (**G**), seedling stage tiller number (**H**), elongation stage tiller number (**I**), flowering stage tiller number (**J**), maturation stage tiller number (**K**), tiller number per plant (**L**), and thousand-grain weight (**M**). Panels (**N-Z**) display heterosis values for these traits (in the same trait order). *P* values indicated independent samples *t*-test or Welch’s *t*-test depending on equality of variances.

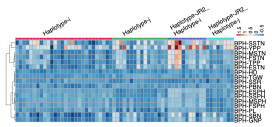

**Figure S3** Heatmap for better-parent heterosis of 17 traits in 63 F_1_ hybrids with homozygous (haplotype-*indica*, haplotype-*japonica*) or heterozygous (haplotype-*indica*_haplotype-JR2, haplotype-*japonica*_haplotype-JR2) haplotypes of *OsBZR1*. Abbreviations: better-parent heterosis = BPH, seedling stage plant height = SSPH, elongation stage plant height = ESPH, flowering stage plant height = FSPH, maturation stage plant height = MSPH, primary branch number = PBN, secondary branch number = SBN, grain number per panicle = GNP, seedling stage tiller number = SSTN, elongation stage tiller number = ESTN, flowering stage tiller number = FSTN, maturation stage tiller number = MSTN, tiller number per plant = TPP, panicle length = PL, seed setting rate = SSR, yield per plant = YPP, heading date = HD, and thousand grain weight = TGW.

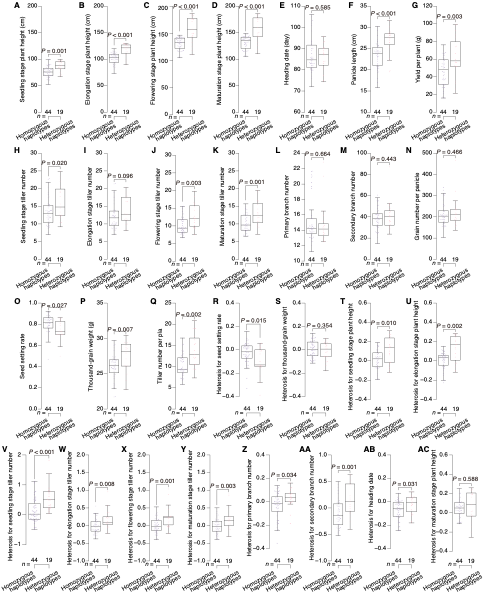

**Figure S4** Phenotypic values and better-parent heterosis of 63 F_1_ hybrids with homozygous (haplotype-*indica*, haplotype-*japonica*) or heterozygous (haplotype-*indica*_haplotype-JR2, haplotype-*japonica*_haplotype-JR2) haplotypes of *OsBZR1*. Panels (**A-Q**) show phenotypic values for seedling stage plant height (**A**), elongation stage plant height (**B**), flowering stage plant height (**C**), maturation stage plant height (**D**), heading date (**E**), panicle length (**F**), yield per plant (**G**), seedling stage tiller number (**H**), elongation stage tiller number (**I**), flowering stage tiller number (**J**), maturation stage tiller number (**K**), primary branch number (**L**), secondary branch number (**M**), grain number per panicle (**N**), seed setting rate (**O**), thousand-grain weight (**P**), and tiller number per plant (**Q**). Panels (**R-AC**) display heterosis values for seed setting rate (**R**), thousand-grain weight (**S**), seedling stage plant height (**T**), elongation stage plant height (**U**), seedling stage tiller number (**V**), elongation stage tiller number (**W**), flowering stage tiller number (**X**), maturation stage tiller number (**Y**), primary branch number (**Z**), secondary branch number (**AA**), heading date (**AB**), and maturation stage plant height (**AC**). *P* values indicated independent samples *t*-test or Welch’s *t*-test depending on equality of variances.

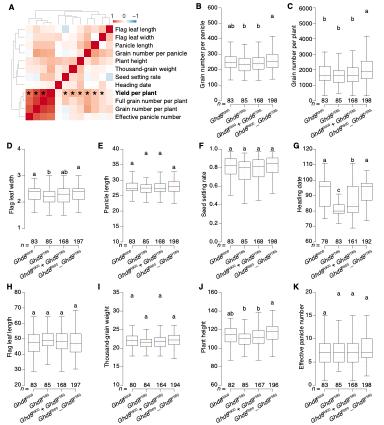

**Figure S5** Phenotypic values of Y900 F_2_ individuals stratified by *Ghd8* genotypes (*Ghd8*^R900^, *Ghd8*^Y58S^, and *Ghd8*^R900^_*Ghd8*^Y58S^), along with group means of homozygous genotypes (*Ghd8*^R900^ + *Ghd8*^Y58S^).  (**A**) Heatmap for correlations among 12 agronomic traits. Panels (**B-K**) show phenotypic values for grain number per panicle (**B**), grain number per plant (**C**), flag leaf width (**D**), panicle length (**E**), seed setting rate (**F**), heading date (**G**), flag leaf length (**H**), thousand-grain weight (**I**), plant height (**J**), and effective panicle number (**K**). *n* = number of F_2_ individuals per group. Analysis of variance with fixed factors in simple linear models was performed to compare the differences, with pairwise comparisons in post hoc test.
